# Adaptive nanopore sequencing for single cell characterization of cancer mutations and gene rearrangements

**DOI:** 10.1101/2022.11.22.517284

**Authors:** Susan M. Grimes, Heon Seok Kim, Sharmili Roy, Anuja Sathe, Carlos I. Ayala, Alison F. Almeda-Notestine, Sarah Haebe, Tanaya Shree, Ronald Levy, Billy T. Lau, Hanlee P. Ji

## Abstract

In this proof-of-concept study, we developed a single cell method that identifies somatic alterations found in coding regions of mRNAs and integrates these mutation genotypes with their matching cell transcriptomes. We used nanopore adaptive sampling on single cell cDNA libraries, generated long read sequences from target gene transcripts and identified coding variants among individual cells. Short-read single cell transcriptomes characterized the cell types with mutations. We delineated CRISPR edits from a cancer cell line. From primary cancer samples, we targeted hundreds of cancer genes, identified somatic coding mutations and a gene rearrangement among individual tumor cells.

## BACKGROUND

Single cell genomics has proven to be a highly informative method for analyzing cancer and other disease tissues. Single cell RNA sequencing (**scRNA-seq**) provides a granular view of an individual cell’s gene expression. One can characterize different cell types, cellular heterogeneity from complex tumor samples and differential gene expression among individual cells. Most scRNA-seq approaches focus on gene expression. However, single cell genomic approaches can examine other features such as copy number and even somatic mutations. These additional genomic features increase the overall yield of valuable information from single cancer cells. However, identifying cancer mutations based on scRNA-seq is not commonly employed given specific limitations of the current short read approach.

The quantitative measurements of single cell mRNA, in the form of sequence reads from the cDNA, requires a combination of cell barcode and a transcript’s sequence. Short read sequencers are used for interrogating scRNA-seq libraries. Short reads are several hundred bases, starting from either the 5’ or 3’ end of a given transcript. This sequence information allows one to count the number of transcripts expressed within an individual cell. However, short reads have significant limitations for the analysis of cancer cell transcriptomes. Many single cell methods require fragmenting the full-length cDNA into lower molecular weight species for preparation of short read sequencing libraries. Fragmenting the cDNA eliminates most of the transcript sequence features that extend beyond the 5’ or 3’ end. This loss includes somatic allelic variants that are present in the internal mRNA coding sequence, chimeric rearrangements and alternative splicing events that alter transcript isoform structure. Overall, short read sequencing leads to a loss of valuable transcript information such as genetic variants that can only be derived from an intact cDNA molecule.

Increasingly, single molecule long read sequencing is used for genomic studies of gene expression and characterizing cDNAs [1–5]. There are two sequencers in this class available from Oxford Nanopore or the Pacific Biosciences. Both generate long reads with lengths from one kb if not higher, all from single DNA molecules. With intact cDNAs from single cells, the library preparation for long read sequencers does not truncate the molecules. As a result, long reads can readily cover an entire cDNA. This sequence information can be used to identify transcript structure and genetic variants present in exon sequences [5].

For single cell genomics, targeted sequencing of specific gene transcripts provides an opportunity to identify transcript structure and genetic variant features present among individual cells. There are a variety of methods used for single cell sequencing of specific target cDNAs. For example, PCR amplification of specific targets from single cell libraries with amplicon sequencing provides higher coverage of genes [6]. Several steps are required for developing PCR assays to amplify gene targets from scRNA-seq libraries. Requirements include identifying specific primer sequences for a given cDNA target and optimizing PCR conditions to reduce the issues of artifacts. When one develops assays for multiplexing PCR, amplification artifacts complicates this method and places practical limits on the total number of amplification targets.

Another common method involves bait capture of cDNAs. These assays use biotinylated oligonucleotide probes which hybridize to a target. This process enriches a specific cDNA molecule of interest from scRNA-seq libraries [2, 3]. The bait capture approach can be scaled up to enrich many genes. However, the development of these assays requires extensive testing of probes and optimizing amplification steps as part of the capture process. The experimental workflow has multiple manipulation steps that add to the complexity of the process.

Several studies have demonstrated a new approach for targeted single molecule sequencing that leverage the attributes of the Oxford Nanopore platform [7]. Referred to as adaptive sampling, this method involves directly assessing DNA molecules for specific target sequences [8–10]. A reference file with a set of the target genomic coordinates is provided. The process involves on-the-fly base calling from each DNA molecule per a given nanopore, sequence alignment of data, real-time control of the nanopore voltage and selection of those molecules with an extended long read of the target sequence. Once the target sequence is identified, the instrument proceeds to sequence the remainder of the molecule. This method enables direct sampling and enrichment of specific DNA molecules without prior preparative steps. It does not require library manipulation to selective PCR amplify or bait hybridization enrichment of the target molecule of interest. Importantly, this approach reduces any potential biases in library content by limiting any pre-amplification step.

We conducted a proof-of-concept study to determine the feasibility of nanopore adaptive sampling applied to scRNA-seq. The objective was to conduct targeted sequencing of specific single cell gene cDNAs and to detect somatic genetic alterations among individual cells from cancer lines or primary tumors. One introduces the single cell cDNA library into the nanopore sequencer - the controller evaluates for matching sequences of a given cDNA molecule as it passes through the pore. Then, the target cDNAs are sequenced with long reads, covering the entire length of the cDNA. Variants that are present in the exons, even when they are positioned far from the 5’ and 3’ ends are detectable. We tested the capability of adaptive sampling for targeting cDNAs from single cell libraries derived from a cancer cell line and identifying mutations or CRISPR edits. Next, we sequenced a set of different cancers including metastatic and lymphoid malignancies. Previously, these patients’ tumors underwent diagnostic cancer gene sequencing – the clinical reports of coding cancer mutation were available for each patient. From these samples and using the same scRNA-seq library, we integrated the single cell RNA-seq short and long read data. The long-read data was used to identify the prior reported cancer mutations and induced CRISPR edits among single cells. We determined if previously reported cancer mutations could be mapped among the single cells from different metastatic sites. Overall, our study demonstrated the feasibility of single cell identification of mutations with adaptive long-read sequencing cDNAs.

## RESULTS

### Overview of the approach

We determined if one could apply nanopore adaptive sampling to single cell detection of somatic genetic variants, mutations and rearrangements (**Figure 1A**). For this study we used a cancer cell line and several tumors that included matched metastases from the same patient. These tumors had prior targeted sequencing results from diagnostic testing. This scRNA-seq approach involved the following steps. We generated single cell cDNAs (10X Genomics) from the sample (**Methods**). A portion of the single cell cDNA undergoes library preparation for conventional Illumina short read sequencing – this type of sequencing requires fragmenting the cDNA. We used an Oxford Nanopore sequencer for providing long reads that span the intact cDNA. Adaptive sampling was used to target sequence-specific genes and their cDNAs. The long reads were pre-processed, aligned and evaluated for CRISPR edits, cancer mutations and rearrangements among the individual cells. Variants were identified from direct examination of the altered position among the sequence reads and variant calling on the long-read data (**Methods**). To infer cell type, we integrated the single cell RNA-seq short and long read data based on matching the cell barcodes. This step allowed us to assign each mutation to specific cell types which is an important step analyzing primary tumor samples. These coding mutations matched those which had already been identified from deep targeted sequencing.

**Figure 1.**
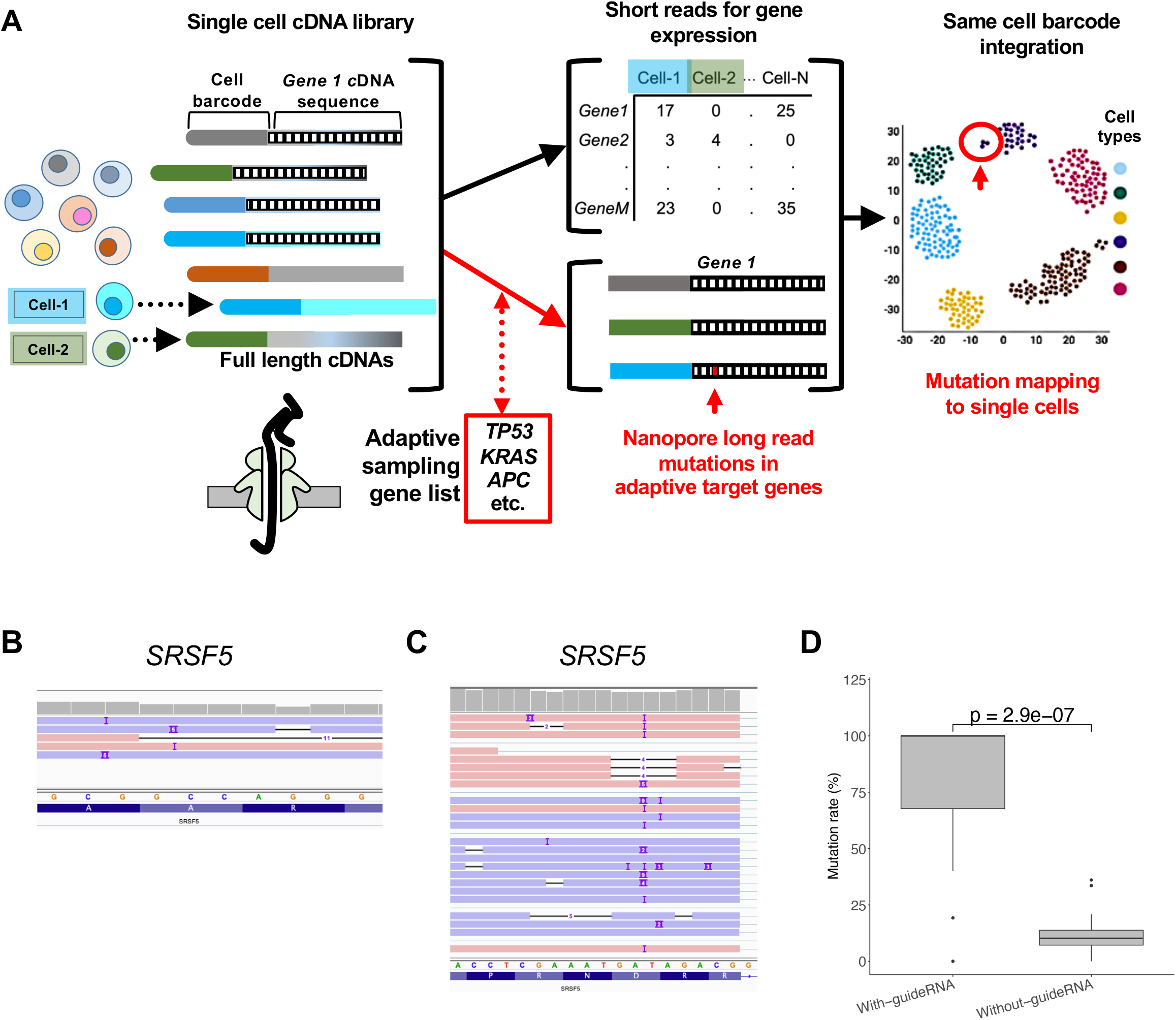
An adaptive sampling methods for sequencing target cDNAs from single cell RNA-seq. A. Overview of single cell library preparation, long and short read sequencing analysis, and integration of results from both modalities. B. IGV screen shot of SRSF5 targeting sites from cells with guide-RNAs: B. SRSF5-1and C. SRSF5-2. D. Boxplot showing CRISPR induced mutation rate for all genes targeted.

### Single cell mutation mapping of a cancer cell line

For the initial experiments, we analyzed the Jurkat cell line which is derived from a T cell leukemia. This cell line has undergone prior genome sequencing with reported mutations [11]. The cells were grown without any CRISPR genome modifications and then underwent single cell cDNA preparation and cDNA amplification. As noted, the same library was split into two aliquots and used for both short and long read sequencing.

We used the results from a short read scRNA-seq analysis of the Jurkat cells to identify individual cells and their gene expression levels (**Methods**). Based on this analysis there was a total of 5,881 cells with an average of 38,966 reads per a cell (**Supplementary Table 1**). Next, we evaluated the expression levels of 319 genes that have been previously reported to have cancer mutations (**Supplementary Table 2**) [11]. These genes covered a range of different expression levels that were corroborated from both the short and long read data (**Supplementary Figure 1**). For example, the *PTGES3L-AARSD1* had the lowest average expression value of 3.4e-04 while the *RPS12* gene had the highest level with a value 5.57e+01.

For mutation identification, we used the second aliquot of single cell cDNA for nanopore adaptive sampling with an Oxford MinION sequencer. For targeting, the adaptive sampling list covered 319 gene targets (**Supplementary Table 2**). The sequencing data was aligned, and the data processed. From the long-read data, there was a total of 5,881 cells (**Supplementary Table 1**). The total number of reads aligning to the target genes were 1.470e+06. The nanopore reads had an average length of 944bp. There was an average of 188 long reads per a cell representing an average of 88.2 genes per a cell. When considering all cells, each gene had an average of 3,733 reads and 1,743 cell barcodes. We compared the cell barcodes between the short and long read data. Short read sequencing had 5,881 cell barcodes of which 5,873 overlapped with the long-read data.

We identified the previously reported mutations (**Methods**). Overall, 292 mutations among 351 mutations were identified among the Jurkat cells, representing 83% of the gene-based mutations that have been previously reported. In total, we identified 910 cells with mutations among 1663 cells. As another general metric for the appearance of a mutation among the cDNA reads, we determined the variant allele frequency per each single cell (**VAF**). This value reflects the ratio of reads identified with the mutation over the total number of reads in a singlecell resolution. The VAF varied among the different genes. For example, the C466Y mutation at *TOP1MT* gene had a mean VAF of 91%, meaning most cells had this mutation. For this mutation, 78 cells had the mutation among 85 cells. In contrast, the VAF for L142L mutation at *ACAT2* gene was 39% (**Supplementary Table 3, Supplementary Figure 2**).

### Single cell mapping of somatic CRISPR edits in Jurkat cells

Next, we assessed whether adaptive sampling and targeted sequencing identified de novo CRISPR-introduced edit mutations from single cells. We used CRISPR-edited Jurkat cells as previously described that stably expressed Cas9 [6]. We transduced this Jurkat cell line with a multiplexed gRNA library containing 32 guide RNAs targeting 16 genes. There were two guides per a gene. We also included five control guide RNAs (**Supplementary Table 4**). These transduced cells underwent processing to generate a single-cell cDNA library. As part of the short-read analysis, we identified which gRNAs were expressed within a given individual cell (**Methods**). This assay relies on using a primer to polymerase extend over the gRNA adjacent to a given cell barcode followed by sequencing in which both gRNA and cell barcode appears in the same read [12]. With the paired gRNA and cell barcode sequence, one determines the distribution of expressed gRNAs across individual cells.

The cells expressing a given gRNA were identified and matched with the long-read adaptive sequencing of single cell cDNAs. With the targeted long reads, we identified the CRISPR-induced edits among the target gene cDNAs among the single cells which also expressed the specific gRNA (**Figures 1B-D and Supplementary Figure 3**). The average number of target long reads matching the gene target was 5.32 per a given cell. As expected, CRISPR mutations were identified at the target gene site among the cells expressing the gRNA. The average target mutation frequency from cells with the guide was 79.0% compared to 11.7% of control group not expressing the gRNA (P = 2.9e–07) (**Figure 1D**).

Furthermore, we detected CRISPR-induced transcript isoform alteration at single cell resolution. For example, the gRNA *SRSF5-2* introduced exon 4 skipping events in 14.81% cells with the guide RNA (**Figures 1B and 1C**). Cells without guide RNA had only 0.43% of those events (**Supplementary Figure 4**). Thus, this result confirmed that adaptive long reads were informative for identifying CRISPR edits.

For validation, we PCR amplified the cDNA targets with long read sequencing. We generated amplicons of the gene targets from the same scRNA-seq library. These amplicons underwent long read sequencing [6]. We identified the matching cell barcodes between the two data sets. Based on comparing the adaptive versus the amplicon sequencing, all identified mutations overlapped, validating the adaptive results (**Supplementary Figure 5**).

### Identifying single cell mutations from tumors

We applied this adaptive nanopore sampling to identify cancer mutations among single cells from tumors. These samples originated from patient biopsies. We analyzed matched pairs of tumors from three patients with metastatic cancer present in different anatomic sites. The first and second patients had appendiceal cancer. The third patient had metastatic follicular lymphoma, a B cell-derived tumor, affecting distinct nodal regions throughout the body.

Every patient had their tumor tested with diagnostic cancer gene sequencing, either from a primary or metastatic site (**Supplementary Table 2**). From the targeted sequencing of each patient’s cancer, we evaluated the reported coding mutations (**Table 2**). The reports only provided nonsynonymous mutations. Using the gene lists from cancer-associated gene panels, we compiled a BED file defining all individual exons and this file was uploaded to the nanopore sequencer workstation for adaptive sampling.

**Table 1.** Cancer samples used for single cell adaptive sequencing.

| Source ID or cell line | Tumor ID | Tumor type | Tumor site or experimental condition | Mutation and somatic variant discovery | Number of coding substitution mutations or CRISPR targets |
| --- | --- | --- | --- | --- | --- |
| Jurkat | C1 | T cell leukemia | Cell line | Exome sequencing | 352 |
|  | C2 |  | Cell line transduced with CRISPR | Amplicon sequencing | 16 (targets) |
| 8605 | T1 | Appendiceal carcinoma | Primary appendix tumor | Cancer sequencing panel | 5 |
|  | T2 |  | Metastasis in the left ovary | Cancer sequencing panel | 5 |
| 8629 | T3 | Appendiceal carcinoma | Metastasis in the omentum | Cancer sequencing panel | 4 |
|  | T4 |  | Metastasis in the small intestine | Cancer sequencing panel | 4 |
| 6408 | T5 | B cell lymphoma | Metastasis in the right inguinal lymph node | Cancer sequencing panel | 9 |
|  | T6 |  | Metastasis in the right cervical lymph node | Cancer sequencing panel | 9 |

**Table 2.**
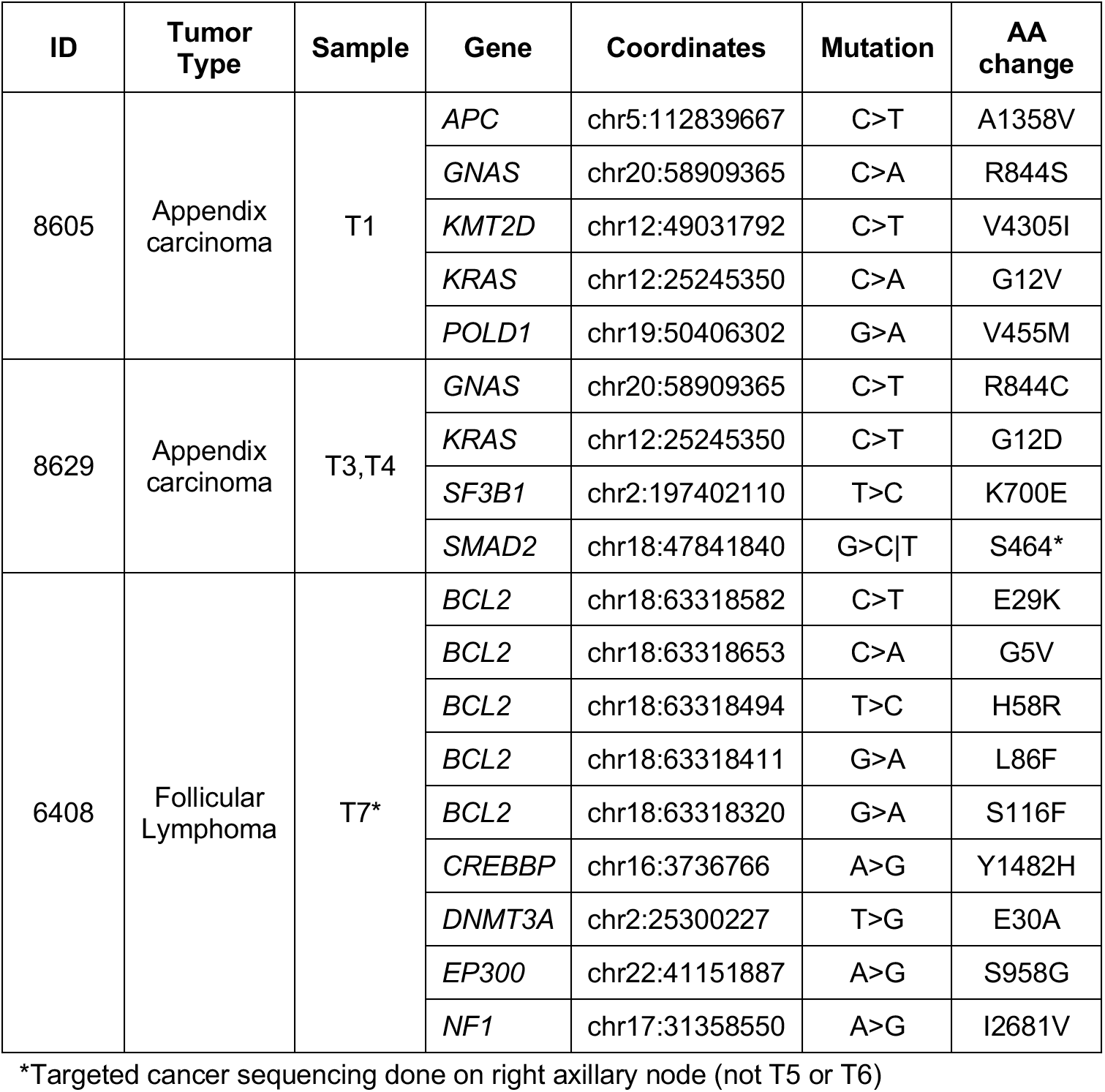
Substitution cancer mutations.

Each tumor sample underwent single cell library preparation and the same single cell libraries were used for both short and long read sequencing (**Methods**). The short-read sequencing provided single cell expression that revealed cell types. The long read alignment provided results for which we identified the base calls at the genomic coordinates of the clinically reported nonsynonymous mutations. This data was also subject to variant calling. Subsequently, we matched the cell barcodes between the long and short read sequences to integrate gene expression, cell type and mutation status.

Across the six tumor samples, we determined the number of single cells by short read sequencing ranged from 7,748 to 16,219 and the median number of genes per cell ranged from 468 to 1,468 (**Supplementary Table 1**). The spread in median genes per cell was attributable to differences in the cell types. Lymphomas are composed of B cells which have a significantly higher number of genes per cell compared to epithelial cells such as those originating from appendiceal cancer. This observation is consistent with what has been noted from single cell studies of lymphocytes and solid tissues [13, 14].

We determined the number of cells by adaptive long read sequencing which had matching cell barcodes compared to the white-list short read data. This ranged between 7,732 to 15,786 per sample (**Supplementary Table 1**). The median transcripts per cell ranged from 10 to 83 and had average number of target genes per cell between 7.6 and 27.0 (**Supplementary Table 1**). Overall, the yield of reads was higher from the B-cell lymphoma than from the appendiceal epithelial tumors.

### Single cell mutations among appendiceal cancers

We analyzed a set of tumors from two patients (P8605 and P8629) with appendiceal cancer. This tumor originates from the epithelial cells of the appendix, a vestigial organ connected to the right colon. The target list covered 529 genes for P8605 and 330 genes for P8629. These lists were based on combining the gene lists for the different sequencing tests. Only coding mutations were reported from these tests.

### 8605’s appendiceal cancer and metastasis

Patient 8605 had an appendiceal carcinoma (**T1**)and a metastatic site (**T2**)located in the left ovary (**Figure 2A**). The patient’s primary tumor site underwent diagnostic cancer sequencing. Based on the clinical report, the T1 tumor had five cancer nonsynonymous mutations in the driver genes *APC*, *GNAS, KRAS, KMT2D* and *POLD1* (**Table 2**).

**Figure 2.**
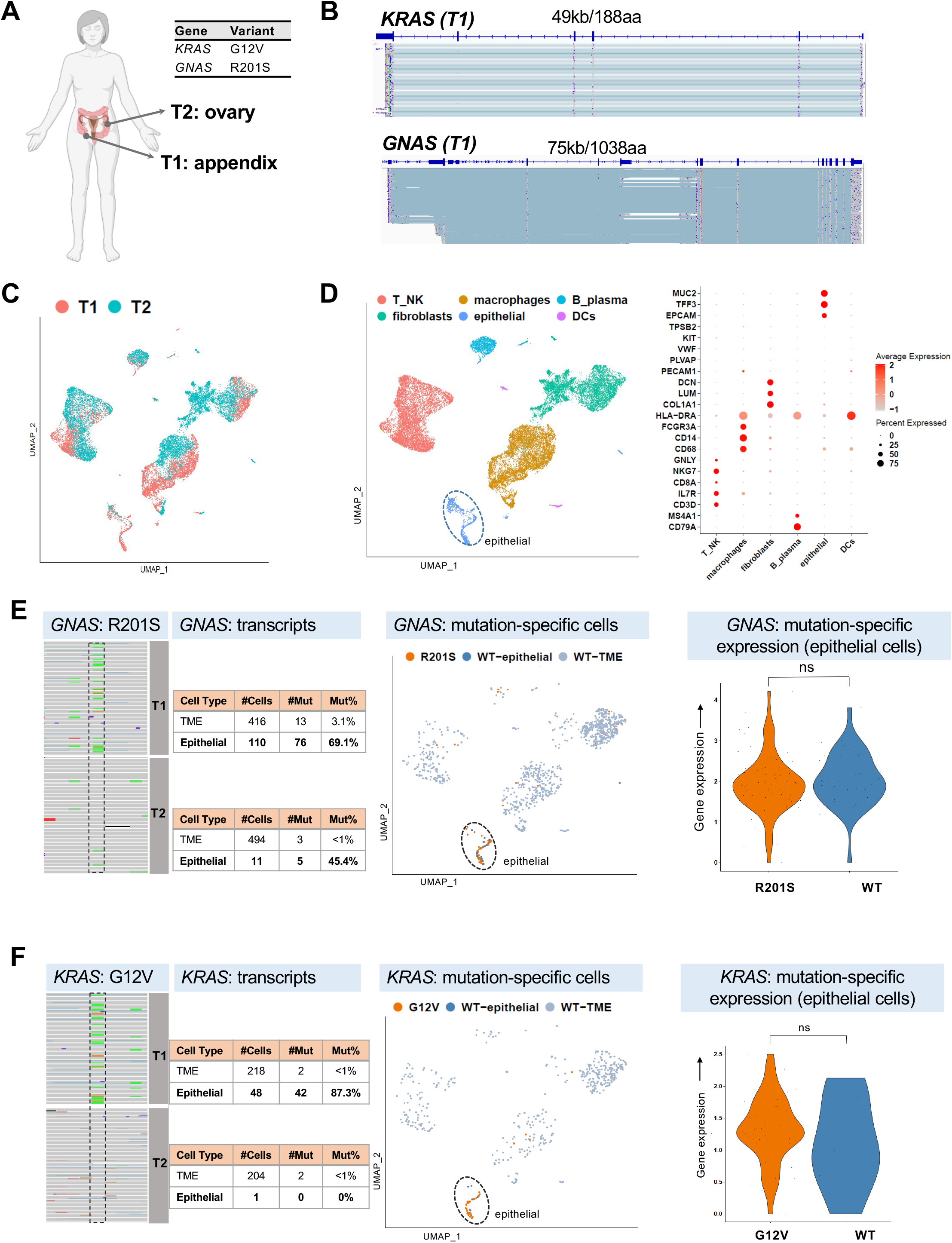
Single cell mutations from the T1 and T2 appendiceal cancers. A. Location of tumor samples for patient 8605, and variants detected from clinical diagnostic sequencing having sufficient long read depth for analysis. B. IGV screen shots for T1 alignments, covering the lengths of *KRAS* and *GNAS* genes. C. UMAP clustered plot showing integration of T1 and T2 samples. D. UMAP clustered plot annotated with cell types, and dot plot showing expression of cell type markers. E. IGV screen shot of T1 and T2 alignments showing *GNAS* R201S mutation position; UMAP plot highlighting location of cells with *GNAS* mutation, and violin plot showing relative expression of mutated and wild type *GNAS* epithelial cells. F. IGV screen shot of T1 and T2 alignments showing *KRAS* G12V mutation position; UMAP plot highlighting location of cells with *KRAS* mutation, and violin plot showing relative expression of mutated and wild type *KRAS* epithelial cells.

The primary appendiceal cancer and its matched metastasis underwent scRNA-seq with both short and long reads (**Figures 2B-D**). The short-read sequencing provided single cell transcriptome information that informed cell identity and the sequencing metrics are shown in **Supplementary Table 1**. Based on the short read sequencing, the T1 appendiceal site had a total of 12,127 cells with an average of 889 genes per a cell (**Figure 2C**). The T2 metastatic site had 14,214 cells with an average of 655 genes per a cell (**Figure 2C**). With this scRNA-seq data, we defined the different cell types in each sample including epithelial, stromal and immune cells. The canonical genes that defined the epithelial cells included *MUC2, TFF3* and *EPCAM* (**Figure 2D**).

Using a sample list of 529 genes, we generated the single cell long-read sequence data from specific target cDNAs. After alignment, we identified the long reads of the target genes which had matching cell barcodes among the short read data (**Methods**). Cell lineages were determined using short-read transcriptome data. We analyzed the adaptive long read data for both tumor sites (**Figures 2E and 2F, Supplementary Table 1**). For the T1 tumor, 67% of long reads matched a short-read barcode: 11,914 cells were identified, along with 498 of the 529 genes targeted. For the T2 metastasis 69% of long read barcodes matched a short read barcode: 14,077 cells were identified along with 496 of the 529 targeted genes. The average number of target genes per cell for the T1 tumor was 11.5 and for the T2 metastasis was 12.8.

As reported from the diagnostic sequencing of the primary tumor, somatic coding mutations were present in *APC*, *KRAS, KMT2D, POLD1* and *GNAS*. All gene mutations were identified in the T1 tumor. The same was true of the T2 tumor, except for *KMT2D* which had no long reads. There were only one or two long-read transcripts for *APC, KMT2D and POLD1* across both samples, limiting the cell type identification for these reads (**Table 3**).

**Table 3.** Single cell identification of cancer mutations.

| Gene | Amino acid change | Tumor ID | Total cells | Wildtype cDNA |  | Mutation cDNA |  | % mut | Tumor ID | Total cells | Wildtype cDNA |  | Mutation cDNA |  | % mut |
| --- | --- | --- | --- | --- | --- | --- | --- | --- | --- | --- | --- | --- | --- | --- | --- |
|  |  |  |  | Epithelial cells | Other cell types | Epithelial cells | Other cell types |  |  |  | Epithelial cells | Other cell types | Epithelial cells | Other cell types |  |
| APC | A1358V | T1 | 2 | 0 | 1 | 0 | 1 | 50% | T2 | 1 | 0 | 0 | 0 | 1 | 100% |
| KRAS | G12V |  | 266 | 6 | 216 | 42 | 2 | 17% |  | 205 | 1 | 202 | 0 | 2 | 1% |
| KMT2D | V4305I |  | 1 | 0 | 0 | 1 | 0 | 100% |  | 0 |  |  |  |  |  |
| POLD1 | V455M |  | 2 | 0 | 0 | 0 | 2 | 100% |  | 2 | 0 | 0 | 0 | 2 | 100% |
| GNAS | R201S |  | 526 | 34 | 403 | 76 | 13 | 17% |  | 505 | 6 | 491 | 5 | 3 | 2% |
| SF3B1 | K700E | T3 | 21 | 6 | 15 | 0 | 0 | 0% | T4 | 9 | 0 | 9 | 0 | 0 | 0% |
| KRAS | G12D |  | 196 | 35 | 131 | 27 | 3 | 15% |  | 236 | 10 | 220 | 5 | 1 | 3% |
| SMAD2 | S464* |  | 33 | 2 | 14 | 17 | 0 | 52% |  | 33 | 0 | 31 | 2 | 0 | 6% |
| GNAS | R844C |  | 322 | 108 | 214 | 0 | 0 | 0% |  | 347 | 17 | 329 | 0 | 1 | 0% |
| GNAS | R844H |  | 382 | 67 | 248 | 59 | 8 | 18% |  | 429 | 12 | 398 | 11 | 8 | 4% |
| Gene | Amino acid change | Tumor ID | Total cells | Wildtype cDNA |  | Mutation cDNA |  | % mut | Tumor ID | Total cells | Wildtype cDNA |  | Mutation cDNA |  | % mut |
|  |  |  |  | B cells | Other cell types | B cells | Other cell types |  |  |  | B cells | Other cell types | B cells | Other cell types |  |
| DNMT3A | E30A | T5 | 57 | 27 | 8 | 19 | 3 | 39% | T6 | 78 | 9 | 35 | 13 | 21 | 44% |
| CREBBP | Y1482H |  | 112 | 59 | 15 | 37 | 1 | 34% |  | 136 | 38 | 61 | 34 | 3 | 27% |
| NF1 | I2681V |  | 29 | 9 | 2 | 14 | 4 | 62% |  | 21 | 7 | 1 | 11 | 2 | 62% |
| BCL2 | S116F |  | 884 | 39 | 8 | 831 | 6 | 95% |  | 620 | 66 | 37 | 497 | 20 | 83% |
| BCL2 | L86F |  | 852 | 43 | 7 | 796 | 6 | 94% |  | 603 | 67 | 37 | 482 | 17 | 83% |
| BCL2 | H58R |  | 756 | 49 | 6 | 696 | 5 | 93% |  | 554 | 24 | 36 | 475 | 19 | 89% |
| BCL2 | E29K |  | 809 | 87 | 7 | 711 | 4 | 88% |  | 574 | 41 | 35 | 479 | 19 | 87% |
| BCL2 | G5V |  | 857 | 844 | 12 | 1 | 0 | 0% |  | 604 | 544 | 59 | 1 | 0 | 0% |
| EP300 | S958G |  | 58 | 37 | 3 | 14 | 4 | 31% |  | 40 | 9 | 10 | 9 | 12 | 53% |

We examined the *GNAS* R201S mutation in the T1 and T2 tumor – for general visualization we combined the data from both tumors for UMAP and violin plots (**Figure 2E**). We determined which T1 and T2 cells had the *GNAS* mutation (**Figure 2E**). *GNAS* is a proto-oncogene that represents the Gsα subunit of heterotrimeric G-proteins and is involved in production of cyclic AMP-based signal transduction [15]. The *GNAS* R201S substitution is a prominent hotspot mutation that results in constitutive activation of the G-stimulatory pathway with an associated increase in cAMP production. For the T1 tumor there were 526 cells with both long and short reads of the *GNAS* gene (**Table 3**). Among T1, the *GNAS* R201S mutation was the most frequently occurring among single cells. We identified 110 epithelial cells: 76 expressed the *GNAS* mutation transcript while 34 expressed the wildtype transcript. For cell types that were not classified as epithelial cells, 403 had the wildtype transcript. Thirteen cells that were not classified as epithelial cells had the mutation – this was likely the result of some variability in the cell assignment using the short read data.

Next, we evaluated the T2 metastasis for this same *GNAS* mutation (**Figure 2E**). There were 505 cells that had both matching long reads of *GNAS* R201S and matching short read transcriptome data (**Table 3**). We identified 11 epithelial cells: five had the *GNAS* mutation transcript while the remaining six expressed the wildtype transcript. Of the remaining 494 non-epithelial cells, over 99% expressed the wildtype *GNAS* transcript.

We identified which cells among the T1 and T2 tumors had the *KRAS* G12V mutation (**Figure 2F**). This mutation is a hotspot that enables *KRAS’s* activity as an oncogenic driver. For T1, there were 266 cells with long reads of the *KRAS* transcript and matching short read transcriptome data (**Table 3**). There were 48 cells that were classified as epithelial: 42 expressed the transcript containing the mutation while 6 expressed the wildtype transcript. Other than epithelial cells, T1 had a total of 218 other cells with the *KRAS* long reads. Two of these cells had the *KRAS* mutations – this may be the result of misclassification. For T2, the *KRAS* G12V mutation had a similar distribution with a total of 205 cells. Two cells had this mutation. The remaining cells all had the wildtype *KRAS* transcript. As noted previously, presence of the mutation suggest that these two cells were epithelial in origin.

There was no evidence of mutation-related nonsense mediated decay in either *GNAS* R201S or *KRAS* G12V epithelial cell transcripts in T1 or T2 (**Figures 2E, 2F**). Transcripts harboring the *GNAS* or *KRAS* mutations had stable gene expression compared to their respective wild type transcripts.

For the T1 tumor, the *KMT2D* V205I and *POLD1* V455M mutations separately mapped to only two cells, and the *APC* A1385V mutation to one cell. In examining the single cell short read data for the T1 tumor, the gene expression levels of *KMT2D, POLD1* and *APC* were generally low with transcript counts between 0.01 and 0.04 per cell (**Supplementary Figure 6**). This low coverage accounts for the reduced number of cells with mutations.

For additional verification of the mutations, we used the Longshot program to call variants from the long read data and confirmed the presence of these mutations (**Supplementary Table 6, Methods**). This program was developed for nanopore long read sequencing [16]. The *KRAS* and *GNAS* mutations were called for T1, and no variants calls were made for *APC*, *KMT2D* or *POLD1*. For the T2 metastasis no mutations were called given the low read depth for *APC*, *KMT2D* and *POLD1* and low variant allele frequency for *KRAS* and *GNAS* (**Table 3**).

### 8629’s appendiceal metastasis

For patient 8629, we had biopsies from two metastatic sites (**T3 and T4**)of an appendiceal cancer (**Figure 3A**). These implants were located on the omentum (**T3**), a tissue covering the abdominal viscera and a metastasis located in the small intestine (**T4**). Based on the diagnostic tumor sequencing, the primary appendiceal tumor had four genes with substitution mutations including *GNAS, KRAS, SMAD2* and *SF3B1* (**Table 2**). These samples underwent both short and long read scRNA-seq (**Figure 3B, Supplementary Table 1**). Based on the short read sRNA-seq, the T3 metastatic site had a total of 10,025 cells with an average of 468 genes per a cell (**Figure 3C**). The T4 metastatic site had 16,219 cells with an average of 578 genes per a cell (**Figure 3C**). With this single cell transcriptome data, we defined the different cell types in each sample including epithelial, stromal and immune cells (**Figure 3D**). After applying standard QC filtering (**Methods**)the T3 site had 8,814 cells of which 1,760 were epithelial, 2,142 stromal and 4,912 were immune cells. The T4 site had 14,511 cells of which 293 were epithelial, 5,863 were stromal and 8,355 were immune cells.

**Figure 3.**
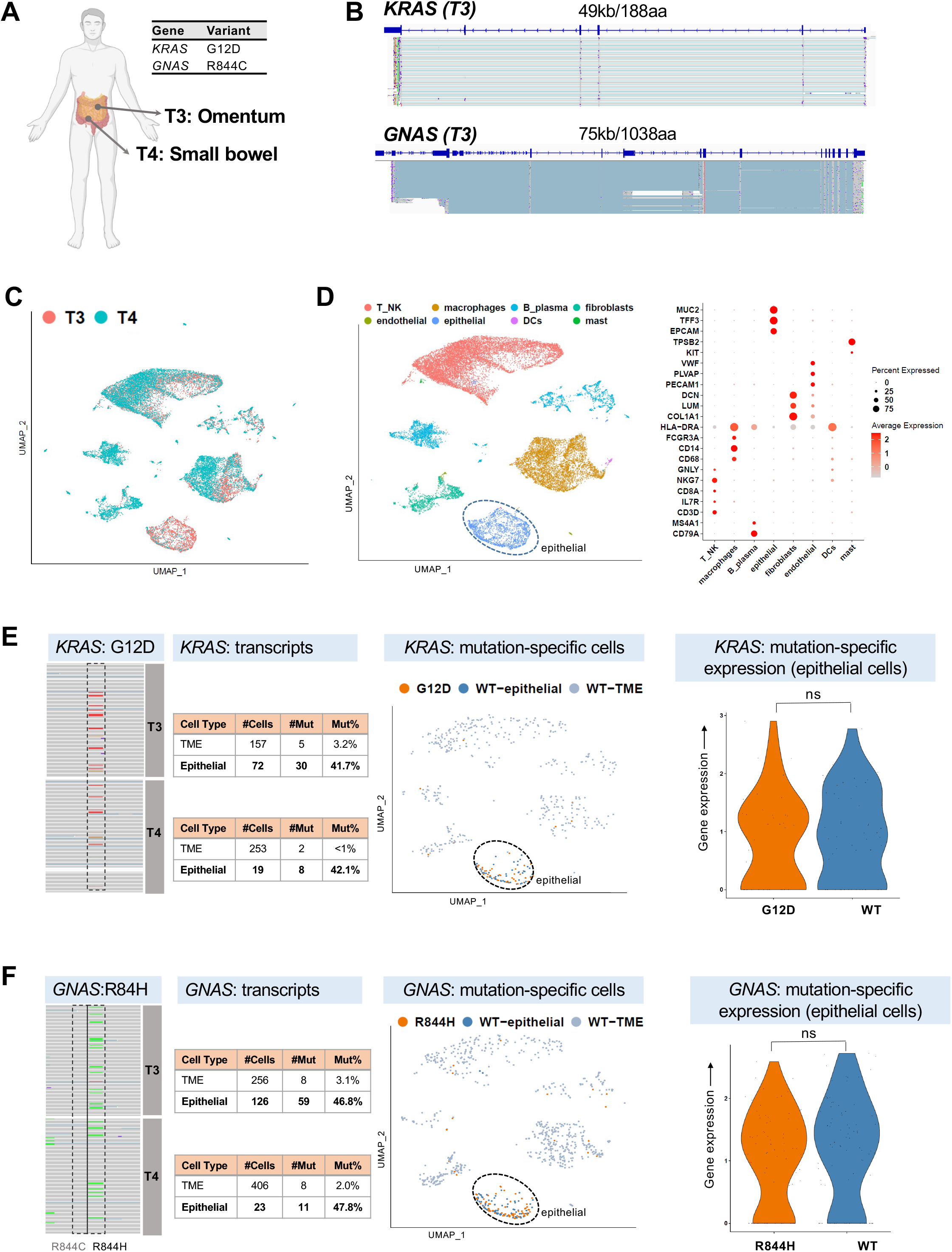
Single cell mutations from the T3 and T4 appendiceal cancers. A. Location of tumor samples for patient 8629, and variants detected from clinical diagnostic sequencing having sufficient long read depth for analysis. B. IGV screen shots for T3 alignments, covering the length of *KRAS* and *GNAS* genes. C. UMAP clustered plot showing integration with T3 and T4 samples. D. UMAP clustered plot annotated with cell types, and dot plot showing expression of cell type markers. E. IGV screen shot of T3 and T4 alignments showing *KRAS* G12V mutation position; UMAP plot highlighting location of cells with *KRAS* mutation, and violin plot showing relative expression of mutated and wild type *KRAS* epithelial cells. F. IGV screen shot of T3 and T4 alignments showing *GNAS* R844C and R844H mutation positions; UMAP plot highlighting location of cells with *GNAS* R844H mutation, and violin plot showing relative expression of mutated and wild type *GNAS* R844H epithelial cells.

We analyzed the long read data for both sites (**Supplementary Table 1**). The target list consisted of 330 genes (**Supplementary Table 2**). For the T3 tumor, 59% of the long read barcodes matched a short read barcode. This data defines a set of 9,929 cells with long read coverage for 312 of the 330 genes. For the T4 site, 70% of long reads matched a short read barcode. This data defined a set of 15,786 cells with long read coverage for 319 of the 330 genes. The average number of target genes per a cell for T1 tumor was 8.4 and the T2 metastasis was 7.6.

There were long reads covering the coding mutation sites for *GNAS, KRAS, SMAD2* and *SF3B* (**Table 3**). For the T3 omental metastasis the *KRAS* G12D mutation was the most prevalent - for general umap and violon plot visualization we combined the data from both T3 and T4 (**Figure 3E**). *KRAS* G12D is a common hotspot mutation found among colon and appendiceal cancers and is a critical oncogenic driver. For the T3 metastasis, 229 cells had both matching long and short read transcriptome data for *KRAS* (**Table 3**). Among this set, there were 72 epithelial cells: 30 had the mutation transcript while 42 were wildtype. For the other cell types, there were total of 157 cells with *KRAS* long reads. These cells included T cells, macrophages, dendritic cells fibroblasts. Among these non-epithelial cells, 152 had the wildtype transcript and five cells had the *KRAS* mutation. Given that these cells had the *KRAS* mutation suggests that they were of epithelial origin. The T4 metastasis had a cellular distribution of the G12D mutation similar to T3 tumor albeit with fewer mutation-bearing cells (**Table 3**). Again, the *KRAS* G12D mutation was the most identified among the single cells. Eight out of the 19 epithelial cells had the mutation (**Figure 3E**).

For T3, the next most frequent mutation found among single cells was the *SMAD2* S464* truncation. This gene is an intracellular signal transducer and transcriptional modulator activated by TGF-beta [17]. There were 33 cells with both long reads and matching short read transcriptome data for *SMAD2* (**Table 3**). Among the T3 epithelial cells, 17 out of 19 cells had long reads mutation transcript. For the non-epithelial cells, there were total of 14 cells, all having the *SMAD4* wildtype transcript. For T2, there were only two epithelial cells, and both had the *SMAD2* S464* mutation. Conversely, the 31 non-epithelial cells had a wildtype SMAD2 transcript.

Finally, for T3 we identified 21 cells with *SF3B1* and 322 cells with *GNAS* (**Figure 3F**). None of these cells had the *SF3B1* K700E or *GNAS* R844C mutations, either among the epithelial cell types or others. However, we did identify a *GNAS* R844H mutation among the T3 cells: this mutation was present among 59 epithelial cells and 8 non-epithelial cells (**Figure 3F**). This finding was notable because it indicated that single cell analysis had a slightly different result about the nature of the mutation – same codon but a different substitution. The difference was likely related to a miscall from the original targeted sequencing report.

The T4 metastasis had a cellular distribution of mutations like the T3 tumor albeit with fewer mutation-bearing cells (**Table 3**). For *GNAS*, 348 cells had the transcript but only one cell had the R844C mutation and is likely to be a sequencing error. Consistent with T3 results, we instead identified a *GNAS* R844H mutation which was present among 11 epithelial cells and 8 non-epithelial cells (**Figure 3F**). Nine cells were identified with the *SF3B1* transcript, but none had the K700E mutation.

There was no evidence of mutation-related nonsense mediated decay in either the *GNAS* R844H or *KRAS* G12D epithelial cell transcripts in T3 or T4. Transcripts harboring the *GNAS* or *KRAS* mutations had stable gene expression compared to their respective wild type transcripts (**Figure 3E, 3F**).

We applied the Longshot variant caller (**Methods**) to identify variants (**Supplementary Table 6**). For the T3 metastasis, the *KRAS* G12D mutation was not identified but the *SMAD2* S464* truncation was called. Of note, Longshot did not identify the *GNAS* R844C but instead identified a *GNAS* R844H mutation, confirming what we noted on evaluation of the mutations in reads among the single cells. This discrepancy in the mutation report likely was an issue with the original variant calling from the first targeted sequencing of the patient’s original tumor. For the T4 metastasis, the mutations were not detected by Longshot, indicating that there were too few cells and read for Long shot to identify the mutation. As already noted, the Longshot caller is not optimized for mutations present at low allelic fractions.

### Single cell mutations and a rearrangement from metastatic B cell lymphoma

For Patient 6408, we analyzed follicular lymphoma samples from two distinct nodal tumor sites (**T5 and T6**). This type of lymphoma is derived from germinal center B-cells and affects the lymphatic system, commonly enlarging the affected lymph nodes. The T5 tumor was obtained from a nodal tumor site in the right groin region and T6 from a nodal tumor site in the right cervical region (**Figure 4A**). The diagnostic sequencing was conducted on a distinct, third lymphoma site from the right axillary lymph node. Coding mutations were reported in five genes that included *BCL2, CREBBP, DNMT3A, EP300* and *NF1* (**Table 2**). All genes have somatic mutations or rearrangements in follicular lymphoma [18].

**Figure 4.**
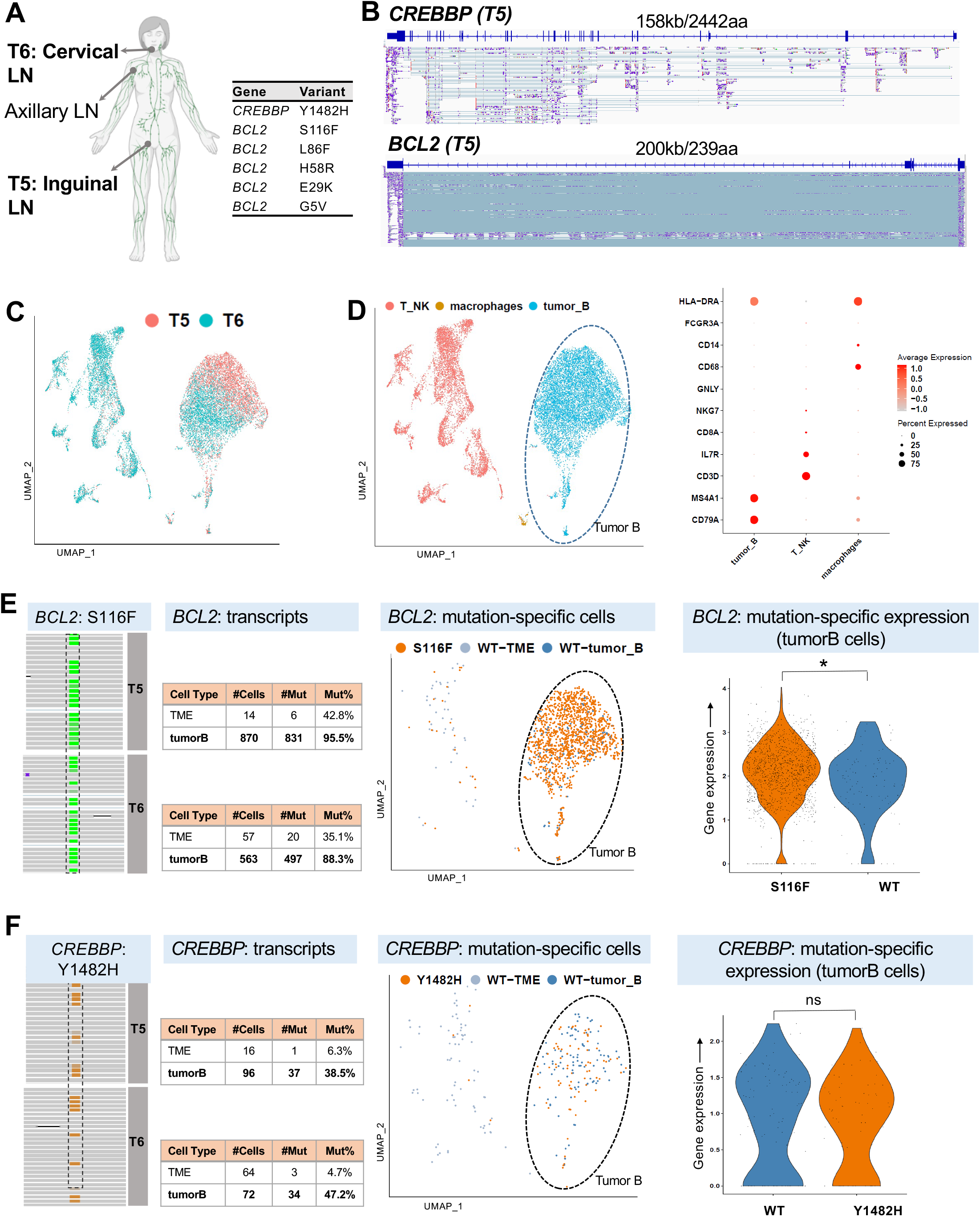
Single cell mutations from the T5 and T6 B cell lymphoma. A. Location of tumor samples for patient 6408 and location of biopsy taken for clinical diagnostic sequencing, plus mutations detected from targeted sequencing having sufficient long read depth for analysis. B. IGV screen shots for T5 alignments, covering the length of *CREBBP* and *BCL2* genes. C. UMAP clustered plot showing integration of T5 and T6 samples. D. UMAP clustered plot annotated with cell types, and dot plot showing expression of cell type markers. E. IGV screen shot of T5 and T6 alignments showing *BCL2* mutation positions; UMAP plot highlighting location of cells with *BCL2* S116F mutation, and violin plot showing relative expression of mutated and wild type *BCL2* S116F B cells, with * indicating significant difference in expression (adjusted p-value < 0.05). F. IGV screen shot of T5 and T6 alignments showing *CREBBP* Y1482H mutation position; UMAP plot highlighting location of cells with *CREBBP* mutation, and violin plot showing relative expression of mutated and wild type *CREBBP* B cells.

For this analysis, the two lymphoma sites underwent both adaptive long and short read scRNA-seq data (**Figure 4A-B, Supplementary Table 1**). Based on the short read sequencing, the T5 site had a total of 7,748 cells with an average of 1,468 genes per cell (**Figure 4C**). The T6 site had 11,865 cells with an average of 1,438 genes per cell (**Figure 4C**). Tumor B cells were identified by variable expression of the immunoglobulin chain as we have previously published [13]. These tumor B cells clustered separately from the macrophages, natural killer and other T cells (**Figure 4D**).

For adaptive sampling of the lymphomas, the target list consisted of 161 genes involved in blood-based malignancies (**Supplementary Table 2**). We analyzed the adaptive long read data for both sites (**Supplementary Table 1**). After matching the cell barcodes between the long and short read data, the T5 tumor had 7,732 cells while the T6 tumor had 11,835 cells. Among the 161 genes from adaptive sampling, we identified 154 and 155 genes for T5 and T6 respectively. The average number of target genes per a cell for T5 right inguinal lymph node was 27 and the T6 right cervical node was 22. Mutations in *BCL2, CREBBP, DNMT3A, EP300* and *NF1* were present among single cells of these tumors with an average of one read with a mutation per a cell (**Figure 4E,4F**). The mutation distribution between the two sites was similar: this means that the relative number of tumor cells with a mutation in each sample were similar across the different genes.

Mutations in *BCL2* were the most prevalent among single cells across both tumor sites (**Table 3 and Figure 4E**). BCL2 inhibits apoptosis and its overexpression prevents cancer cell death [18]. *BCL2* is typically overexpressed in follicular lymphoma due to a hallmark t(14;18)(q32;q21) *IGH/BCL2* translocation ir enhancer sequences of th chromosome 14 [18]. In t hypermutation. This high *IGH/BCL2* translocation involving the BCL2 gene on chromosome 18 translocated to the enhancer sequences of the immunoglobulin heavy chain gene (*IGH*) promoter region on chromosome 14 [18]. In the presence of this translocation, *BCL2* is a target of somatic hypermutation. This high mutation rate is a result of activation-induced cytidine deaminase activity which alters cytosine in DNA. *BCL2* mutations in T5 and T6 were clustered in a hotspot and were all phased, meaning they were ordered tandemly on the same molecule, representing a somatic mutation haplotype. This clustering of *BCL2* mutations was reported from the targeted sequencing of the third tumor site. Our adaptive sampling results confirmed this finding for four of the five *BCL2* mutations, with between 87% and 94% of cells harboring each of the four mutations. The fifth *BCL2* mutation was observed in only one of 844 cells spanning that genomic location.

In the T5 tumor, the next most frequent mutation was *CREBBP* Y1482H (**Table 3 and Figure 4F**). This transcript was observed among a total of 112 cells. The mutation was found among 37 of the 96 tumor B cells and in one of the other 16 cells in the TME. The *EP300* S958G, *DNMT3A* E30A and *NF1* I2681V mutations were found at between 31% and 53% frequency among the tumor B cells (**Table 3**). As noted, the T6 metastasis had a mutation pattern like T5 tumor albeit with fewer cells and in general a similar or slightly lower percentage of mutation-bearing cells (**Table 3**).

There was no evidence of mutation-related nonsense-mediated decay in *CREBBP* Y1482H B-cell transcripts in T5 or T6. Transcripts harboring the *CREBBP* mutation had stable gene expression compared to the wildtype transcript (**Figure 4E**). *BCL2* gene expression in tumor B-cells was moderately higher in transcripts harboring the S116F mutation compared to the wildtype transcript, with p-value 0.011 using Welch two sample t-test (**Figure 4F**).

As an additional verification, we used Longshot to identify mutations from the long read data. Four of the five *BCL2* mutations were called in the T5 lesion, as well as the mutations in *DNMT3A, CREBBP, NF1* and *EP300*. Read depth at the fifth *BCL2* mutation was high as with the other four mutations. However, there was no mutation present at this position. In the region between the first and fourth *BCL2* mutation three other variants were called by Longshot and supported by visual inspection of the reads (**Figure 5A**). This result is consistent with a somatic hypermutation event in *BCL2*. The positive and negative mutation calls for T6 were identical to T5: four of the five *BCL2* mutations were called, plus the mutations in in *DNMT3A, CREBBP, NF1* and *EP300* (**Table 3**). In contrast however to T5, only two of the three additional *BCL2* variants were found. The variant at chr18:63318573 which was heterozygous in T5, did not appear in T6.

**Figure 5.**
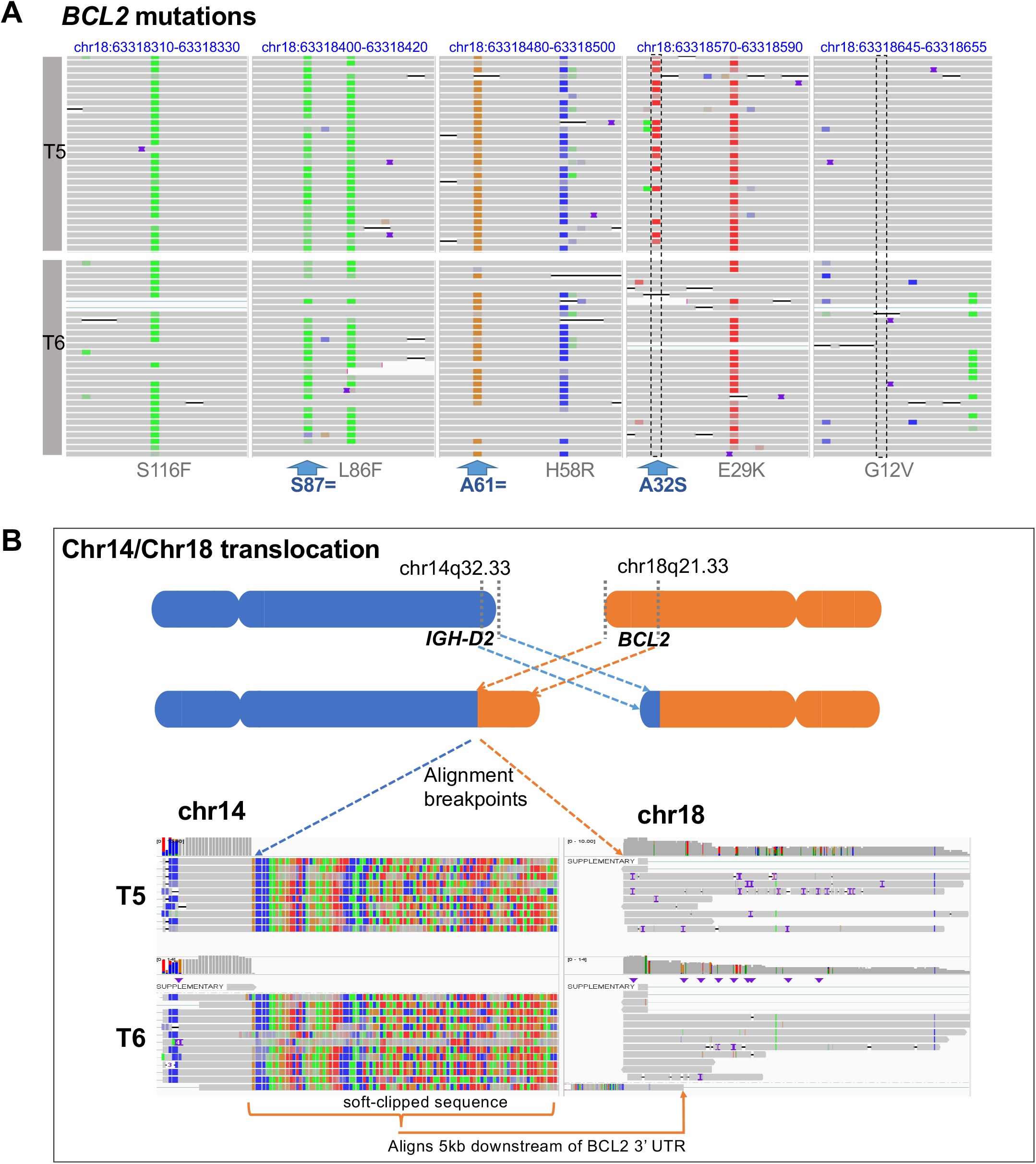
A. IGV screen shots of T5 and T6 lymphomas and locations of *BCL2* variants called by Longshot. Coding mutations are labeled in grey and additional variants detected by Longshot labeled in blue. B. Schematic of translocation detected by cuteSV. An IGV screen shot showing primary alignments to *IGH-D2* on chromosome 14 with soft-clipped sequence to the right, plus secondary alignments of the same reads to a region downstream of *BCL2* 3’UTR on chromosome 18.

Finally, we used the cuteSV program to call structural variants from T5 and T6 [19]. An *IGH/BCL2* rearrangement was identified in both the T5 and T6 tumors (**Figure 5B**). The breakpoints were in *IGH-D2* and approximately 5kb downstream from *BCL2*3’ UTR. We determined that multiple long reads supported the rearrangement. This result represents the first demonstration where single cell sequencing reveals the presence of a rearrangement chimeric transcript.

## DISCUSSION

This proof-of-concept study demonstrates a new approach for single cell identification of cancer mutations. This method integrates nanopore adaptive sequencing and single cell RNA-seq.

With the adaptive sampling feature of Oxford Nanopore’s sequencer, one selects specific target cDNAs, derived from mRNAs, based on a list of gene coordinates. Our largest gene list consisted of 529 genes. Adaptive sampling enabled these targets to be sequenced with an enriched number of reads compared to the remainder of the cDNA population. The nanopore long reads cover the entire cDNA which enables one to determine if coding mutations are present in the mRNA sequence. The same single cell cDNA library is also applied to conventional short read sequencing which provides the transcriptome features of the same cells. By matching cell barcodes, the long and short read data are integrated, thus providing both full length mRNA sequence features and single cell gene transcriptomes. We applied this targeting feature to single cell cDNA libraries generated from a cancer cell line and tumor biopsies. These samples had undergone prior genome sequencing of cancer genes and had lists of the coding mutation that result in substitutions. Overall, we identified these coding mutations among single cells from the cancers. Likewise, one of the cancers had a translocation leading to a chimeric transcript. This rearranged cDNA was present and identified among single cancer cells. This results demonstrates a potentially useful approach for identifying gene chimeras resulting from translocations and other types of rearrangements.

Targeted single cell RNA-seq has advantages related to lower cost compared to single cell whole transcriptomes and higher read coverage. The adaptive sampling method in scRNA-seq offers these advantages and others. Importantly, integrating this targeted long read approach provides a new way to increase the yield of genomic features from single cell gene expression studies. For this proof-of-concept, all that was required was retaining a portion of the single cell cDNA library prior to the fragmentation for short read library preparation. Another advantage comes from the workflow. Since the target DNA molecules are selected for sequencing based on their sequence properties, there is no need for any prior enrichment steps. In addition, cDNA fragmentation is not required since long read libraries are compatible with full length cDNA. Overall, this reduces the complexity of library preparation.

This approach provides a way to identify CRISPR edits and enables one to screen for CRISPR genotypes directly so long as they occur within coding regions. Some CRISPR edits may lead to nonsense mediated decay. This change in gene expression may also be detected by this method based on comparing the levels of gene expression with wildtype cells.

This study identified specific issues of adaptive sampling for identifying transcript-based mutations with scRNA-seq. Because the sampling depends on the intrinsic expression levels of a given mRNA / cDNA, transcripts with low expression will provide fewer molecules for sequencing. When analyzing single cells, the transcript yield is already low. Therefore, some transcripts with low expression are missed and this reduces this single cell representation. The sensitivity of detecting mutations from low abundance transcripts is reduced. One approach to overcome this limitation involves enriching and amplifying the target genes from a single cell cDNA library. Our future work will involve integrating adaptive nanopore sampling and single cDNA targeting.

There are many potential applications for this approach. For example, one could identify specific different mutations that define the subclonal populations of a tumor. This type of analysis may prove useful in the study of other diseases beyond cancer. For example, clonal hematopoiesis of indeterminate potential involves hematopoietic stem cells which have genetically distinct subpopulations defined by the presence of somatic mutations. This approach provides a way to determine which cell types account for mutations with low allelic fractions that were identified with bulk genomic DNA sequencing. As we demonstrated, this approach can identify gene fusions and may provide a new way of screening cancers for rearrangements. As we have described in our previous work, targeted sequencing of specific cDNAs provides detailed information about transcript isoforms which play a key role in regulating cell terminal differentiation. Thus, one could have integrated long and short read analysis to define the associations of alternative isoforms in specific cell types.

## CONCLUSION

In this proof-of-concept study, we developed a single cell method that identifies somatic alterations found in coding regions of mRNAs and integrates these mutation genotypes with their matching cell transcriptomes. Based on nanopore sequencing and adaptive sampling of single cell cDNA libraries, we identified CRISPR edits, somatic mutations and gene rearrangement at single cell resolution. Combining this genotype information with single cell gene expression allows us to infer which cells had these somatic alterations.

## METHODS

### Patient samples and processing

Patients with metastatic appendiceal cancer were consented with an IRB protocol 44036 approved by Stanford University. The patient with FL was consented with an IRB protocol 36750 approved by Stanford University. Fine needle aspirate specimens from two spatially separated nodal tumor sites were obtained and subjected to scRNA-seq and additional targeted DNA sequencing was performed on separate banked formalin-fixed paraffin-embedded lymphoma biopsies from a third tumor site. Excisional biopsies from tumor tissue were obtained from surgery and stored in RPMI medium before dissociation. Single-cell suspensions were obtained from tissue fragments using enzymatic and mechanical dissociation. Cells were washed twice in RPMI + 10% FBS, filtered through 70 μm (Flowmi, Bel-Art SP Scienceware, Wayne, NJ), followed by 30 μm (Miltenyi) or 40 μm filter (Flowmi). Cryofrozen cells were rapidly thawed in a bead bath at 37 °C followed by above washing and filtering steps. Live cell counts were obtained on a BioRad TC20 cell counter (BioRad, Hercules, CA) or a Countess II FL Automated Cell Counter (ThermoFisher Scientific) using 1:1 trypan blue dilution. Cells were concentrated between 500-1500 live cells/μl for subsequent single cell library preparation.

### Cell lines and induction of CRISPR mutations

The Jurkat cell line (ATCC TIB-152) and a Cas9-stable version of Jurkat (SL555, GeneCopoeia, Inc., Rockville, MD, USA) were maintained in RPMI medium supplemented with 10% FBS at 37C under standard CO2 conditions. We produced an oligonucleotide pool for the gRNA library (IDT, Coralville, Iowa, USA). Amplified gRNAs were cloned to lentiGuide-Puro (Addgene plasmid #52963). To transduce Cas9-expressing Jurkat cell for CRISPR editing, we used the spinoculation method. The lentiviral supernatant and 8μg of polybrene (Sigma-Aldrich, MO, USA) were added to 1.0 × 10^5^ Cas9-stable Jurkat. The mixture was centrifuged at 800g at 32 Celsius degree for 30 minutes. Cell pellets were resuspended to fresh media and after 72 hours, transduced cells were selected by puromycin (Life Technologies, CA, USA). Additional details about this CRISPR edited cell line are fully described by Kim et al. [6].

To identify specific CRISPR mutations, we generated single cell full length cDNAs from transduced Jurkat cells as previously described [6]. One ng of single cell cDNA library was used to amplify transcripts with a set of primers flanking the CRISPR edit site. KAPA HiFi HotStart ReadyMix (Roche, Basel, Switzerland) was used for amplification. Extension time was 60s. Amplicons were pooled at equimolar concentrations. The libraries were prepared with 900fmol of pooled amplicon for Promethion flow cell FLO-PRO002 (Oxford Nanopore Technologies) using Native Barcoding Expansion and Ligation Sequencing Kit (Oxford Nanopore Technologies) as per the manufacturer’s protocol. Libraries were sequenced on Oxford Nanopore Promethion for 72h.

### Single cell library preparation and short read sequencing

Sequencing libraries were prepared using Chromium NextGEM Single Cell 5’ Library & Gel Bead Kit v1.1 or v2 (10X Genomics, Pleasanton, CA, USA) as per the manufacturer’s protocol. Guide RNA direct capture for Jurkat CRISPR assay has been performed as previously described using 6pmol of scaffold binding oligo nucleotides [6]. The cDNA and gene expression libraries were amplified with either 14 or 16 cycles of PCR, depending on the starting amount. The size distribution of gene expression libraries was confirmed via gel electrophoresis (ThermoFisher Scientific, Waltham, MA, USA). The libraries were quantified using a Qubit fluorescent assay (Invitrogen). Short-read sequencing was performed on Illumina sequencers (Illumina, San Diego, CA, USA).

### Nanopore long read sequencing of single-cell libraries

We amplified the entire single-cell cDNA material using the following primer sequences: Partial Read 1: CTACACGACGCTCTTCCGATCT and Non-polydT: AAGCAGTGGTATCAACGCAGAG. KAPA HiFi HotStart 2X ReadyMix (Roche, Basel, Switzerland) was used for PCR amplification with 250 nM of each primer. Following PCR, the amplicons were purified using 1.5X volume equivalents of Ampure XP beads. Libraries were quantified with Qubit (Thermo Fisher Scientific). The library was diluted to a total concentration of 600 fmol prior and loaded onto a MinION R9.4.1 flow cell and sequenced for 72 hours per the manufacturer’s instructions (LSK-110, Oxford Nanopore Technologies). For the Promethion runs, 900fmol of pooled amplicon were loaded onto a Promethion flow cell FLO-PRO002 (Oxford Nanopore Technologies) and sequenced for 72 hours.

Each patient had one their tumor sites analyzed with three different cancer gene panels used for diagnostic tumor sequencing. Among these different tests, the number of target genes ranged from 130 to 529. We merged these gene lists based on the type of test that conducted on the patient’s tumor. Our samples included hematologic and solid epithelial tumors. For each patient, we had a list of the reported somatic mutations. From the merged gene lists, the exon locations were identified and organized into a bed file. For adaptive sequencing, we uploaded this genomic bed file into the instrument control software. Live basecalling was based on the ‘fast’ model enabled rapid alignment and subsequent enrichment of reads that overlapped the target regions.

### Bioinformatic analysis

#### Short-read processing and cell type assignment

Cellranger (10x Genomics) version 3.1.0 ‘mkfastq’ and ‘count’ commands were used with default parameters and alignment to GRCh38 to generate matrix of unique molecular identifier (UMI) counts per gene and associated cell barcode. We constructed Seurat objects from each dataset using Seurat (version 4.0.1) [20, 21] to apply quality control filters. Quality controls included removing cells that expressed fewer than 200 genes, had greater than 30% mitochondrial genes or had UMI counts greater than 6000 indicating potential doublets. If a gene was detected in less than three cells it was removed. We normalized data using ‘SCTransform’ and used first 20 principal components with a resolution of 0.8 for clustering. We then removed computationally identified doublets from each dataset using DoubletFinder (version 2.0.2) [22]. The ‘pN’ value was set to default value of 0.25 as the proportion of artificial doublets. The ‘pK’ value representing the PC neighborhood size was calculated using 20 principal components. pK’ value representing the PC neighborhood size was calculated using 20 principal components. The ‘nExP’ value was set to expected doublet rate according to Chromium Single Cell 3’ v2 reagents kits user guide (10X Genomics). These parameters were used as input to the ‘doubletFinder_v3’ function to identify doublet cells.

For determining cell type, clusters were annotated based on canonical known genes markers. Among our tumor biopsies, we had appendiceal carcinomas which are epithelial in origin and lymphomas which are B cell derived. For the appendiceal cancers, we identified epithelial cells (*EPCAM, TFF3, MUC2*), fibroblasts (*DCN*, *COL1A1, LUM*), endothelial cells (*VWF, PLVAP, PECAM1*), T cells (*CD3D*, *IL7R, CD8A*), NK cells (*NKG7, GNLY*), B or plasma cells (*MS4A1*, CD79A), mast cells (*TPSAB1*) and macrophages or dendritic cells lineages (*CD68, CD14, FCGR3A, HLA-DRA*).

For the lymphoma samples, we included *MS4A1, CD19, CD79A* (B cells), *CD3E, CD3D, CD2* (T cells), *CD8A, CD8B* (CD8+ T cells), *CD4* (CD4+ T cells), *LEF1, CCR7, NOSIP* (Naive T cells), *IL7R, SELL* (Memory T cells), *CD4*, *IL2RA, FOXP3* (T regulatory cells), GZMA, NKG7 (T effector cells), GNLY, NCAM1 (NKT/NK cells), and *CD14, LYZ* (myeloid cells). As a cross reference, we validated our results with cluster markers with previously characterized gene expression profiles of sorted cell types that included lymphocytes. In the case of the lymphomas, the classification of malignant versus non-malignant B-cell cells was based on calculating the average expression of each kappa and lambda variable region gene for the different clusters. The expression of a clonal light chain provided assignment for the malignant B cell clusters. In contrast, the normal B cell cluster expressed heterogeneous BCR light chain variable genes.

#### Adaptive long read processing

The adaptive sequencing runs from the Oxford Nanopore and their sequence output was filtered to include just the reads within one of the targeted regions. This step involved using the log file provided by the sequencer. The log file indicates whether each read was ejected (‘unblock’) or accepted (‘stop_receiving’ – enriched). The accepted reads, which contain full length cDNA, were bioinformatically selected using the fast5_subset command from the ont_fast5_api package. This data was iteratively processed using the ‘super-accuracy’ basecalling mode with Guppy (v5.0.16) and was aligned to the reference genome GRCh38 using minimap2 (v2.22) [23].

#### Integration of short and long-reads from single cell cDNA

As previously described, we developed a method to match the short and long reads from overlapping single cells [6]. We compared a whitelist of cell barcodes identified in the short reads with barcode sequences extracted from the soft-clipped sequences in the aligned long reads. The python pysam module was used to identify soft-clipped portions of aligned reads. The next step was a machine learning approach utilizing a cosine-similarity function (CountVectorizer from scikit-learn python module, with kmer length of 8) to identify potential barcode matches within the soft-clipped sequences. Using the five highest ranking cosine similarity scores per read, the edit distance between the long read barcode sequence and the whitelisted barcode was calculated. Barcode matches with the lowest edit distance. In cases where there was a tie, the highest cosine similarity score was selected for final evaluation. If the paired barcode edit distance was less than 3 it was considered a successful match, otherwise the read was not considered a match to any of the short-read barcodes and was excluded from further integrated analysis. From the resulting file, any exactly matched barcode/UMI combinations were removed as PCR duplicates.

### CRISPR genotyping analysis

Using the long read data from targeted cDNAs, we identified the genotypes of CRISPR mutations from the Jurkat cell line. After aligning each nanopore read and confirming the coordinates of the target, we evaluated a 2bp sliding window that was tiled across the putative cleavage site. Insertions, deletions or base substitutions were identified among the long reads.

We performed this analysis for each gRNA target per a given cell and summarized the mutation frequency of the CRISPR target.

### Single cell analysis of cancer mutations from tumor biopsies

We had a set of tumor samples originating from patients with metastatic cancer. These patients had one of their tumor sites undergo diagnostic cancer genome sequencing. The clinical sequencing reports provided a list of mutations which led to amino-acid changes, frameshifts or premature stops – this information was compiled for our study. For mutations reported in GRCh37 coordinates, we conducted a liftover procedure to convert to GRCh38 coordinates.

For mutations reported as amino-acid changes, we conducted an analysis with the CADD application to lift these mutations to the GRCh38 reference coordinates [24]. We used the pileup command from the python pysam module to identify the specific nanopore reads which had the reported mutations [25, 26]. As an additional validation of the tumor mutation calls, we used Longshot to call variants [16]. Longshot is designed for germline variant calling of long reads, so some parameters were adjusted to provide more sensitive variant calling appropriate for somatic mutations. Longshot was run with variant phasing disabled, a strand bias p-value cutoff of 0.0001, and variant density filter set to 10:100:50 (10 variants within 100 base pairs with genotype quality >=50) to filter out any variants in a very dense cluster. The cell barcode for each long read was identified as described in the paragraph on adaptive long read processing above and using this information the cells in the short-read Seurat object were annotated as having either reference or alternate base values. Standard Seurat functions such as DimPlot and VlnPlot were then used to visualize the differences in cell type distribution and gene expression level, for those cells with the mutation versus the wildtype.

To determine if there were rearrangements, we used cuteSV [19]. The following parameters were applied: maximum distance to cluster reads together for insertion or deletion: 100, maximum basepair identity to merge breakpoints for insertion or deletion: 0.3.

### Transcript isoform analysis

For each read, we used the exon coordinates for the targeted gene, to determine which exons were present and therefore the isoform structure [6]. Exon coordinates were based on the Ensembl canonical transcripts from the GENCODE version 38 GTF file [27]. To assign exons for a given cDNA, we required that the read had greater than 12 aligned bases from a specific exon. We merged the information about the read mutation status and the isoform structure. This data was summarized to provide isoform counts by mutation status (reference sequence versus alternative allele) for each gene.

## Abbreviations

AMP: Adenosine monophosphate
BED: Browser Extensible Data format
cAMP: Cyclic adenosine monophosphate
CRISPR: Clustered Regularly Interspaced Short Palindromic Repeats
gRNA: guide RNA
QC: Quality Control
scRNA-seq: single-cell RNA sequencing
SNV: Single nucleotide variant
TME: Tumor microenvironment
VAF: Variant Allele Frequency

## DECLARATIONS

### Ethics approval and consent to participate

This study was conducted in compliance with the Helsinki Declaration. The Institutional Review Board at Stanford University School of Medicine approved the study protocols (IRB-11886 and IRB-48492). All patients provided written informed consent to participate.

### Consent for publication

All patients consented for publication of the de-identified results.

### Availability of data and materials

The sequencing data have been deposited in the NCBI Sequence Read Archive database under the accession number PRJNA708300 [23]. Scripts for analysis are publicly available on github (https://github.com/sgtc-stanford/scCRISPR) [24] and zenodo (https://zenodo.org/badge/latestdoi/365008149) [25] under MIT license.

### Competing interests

The authors declare that they have no competing interests.

### Funding

This work was supported by US National Institutes of Health grants R33 CA247700 (HPJ, HSK, BTL, SR), R01HG006137 (HPJ, HSK) and R35HG011292-01 (BTL, HSK). HPJ also received support from the Clayville Foundation. Additional funding came from U.S. National Institutes of Health R35 CA197353 (RL), 5T32HL120824-04 (TS) and K08CA252637 (TS). Other funding came from Leukemia & Lymphoma Society grant TRP 6539-18 (RL), and the Hoogland Lymphoma Research Fund (RL), the American Cancer Society Postdoctoral Fellowship PF-17-239-01-LIB (TS). SH was supported by a Mildred Scheel postdoctoral fellowship from the Deutsche Krebshilfe (70113507).

### Authors’ contributions

SMG, HSK, SR, BTL and HPJ were involved in conception and design of the study, development of methodology, acquisition of sequencing data, analysis and interpretation of data and writing of the manuscript. AAP and CIA managed the acquisition of samples. SH, TS and RL provided lymphoma samples and single cell libraries. SMG, HSK and HPJ were involved in analysis and interpretation of data. SMG, HSK, BTL and HPJ were involved in the development of analysis methodology and acquisition of data. SMG, HSK, SR and HPJ wrote the manuscript. HPJ oversaw all aspects of the study.

## Acknowledgements

Not applicable.

## ADDITIONAL FILES

**Additional file 1: Supplementary Figures**

**Fig S1**. Jurkat wild-type long-read and short-read gene expression.

**Fig S2**. IGV screenshot of genetic mutations in Jurkat cell-lines.

**Fig S3**. Single-cell level quantification of CRISPR induced indel per each guide-RNA.

**Fig S4**. Single-cell level quantification of CRISPR induced exon skipping.

**Fig S5**. Comparison of single-cell level quantification of CRISPR induced indel between amplicon vs adaptive sequencing.

**Fig S6**. Comparison of long-read and short-read gene expression for patient tumor samples

**Additional file 2: Supplementary Tables**

**Table S1**. Sequencing metrics.

**Table S2**. List of target genes for nanopore adaptive sequencing.

**Table S3**. Jurkat mutations.

**Table S4**. Guide RNA sequences.

**Table S5**. CRISPR primer list.

**Table S6**. Longshot variant calls proximal to clinical diagnostic mutations.

## Notes

### Competing Interest Statement

The authors have declared no competing interest.

