## Supplemental Figures for "Adaptive nanopore sequencing for single cell characterization of cancer mutations and gene rearrangements"

**Supplementary Figure S1. Jurkat wild-type long-read and short-read gene expression.** This graph shows the single cell expression level of each gene analyzed by short-read sequencing (x-axis) versus long-read sequencing (y-axis). Gene expression from short-read sequencing is analyzed in Seurat and long-read sequencing by unique molecular index counting.

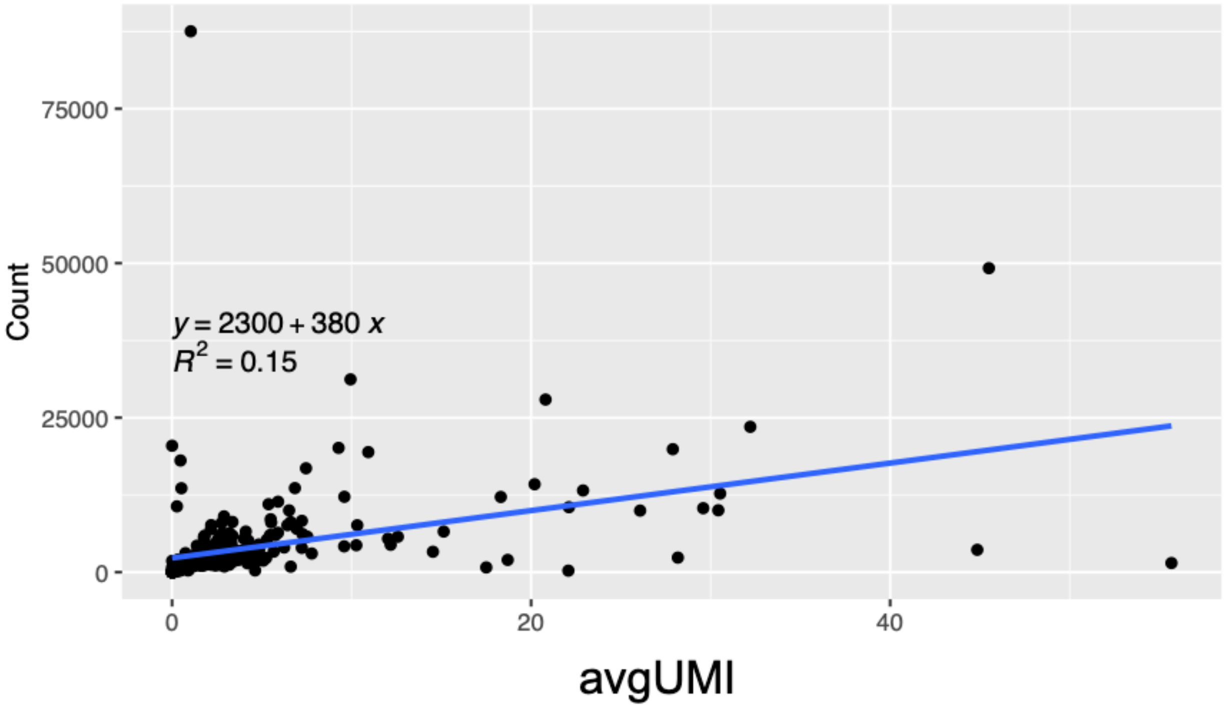

**Supplementary Figure S2. IGV screenshot of genetic mutations in Jurkat cell-lines from long reads.** Sequence reads from the Jurkat cell single-cell cDNA spanning *TOP1MT* (upper panel) and *ACAT2* (lower panel). Mutations were identified at expected positions as previously reported.

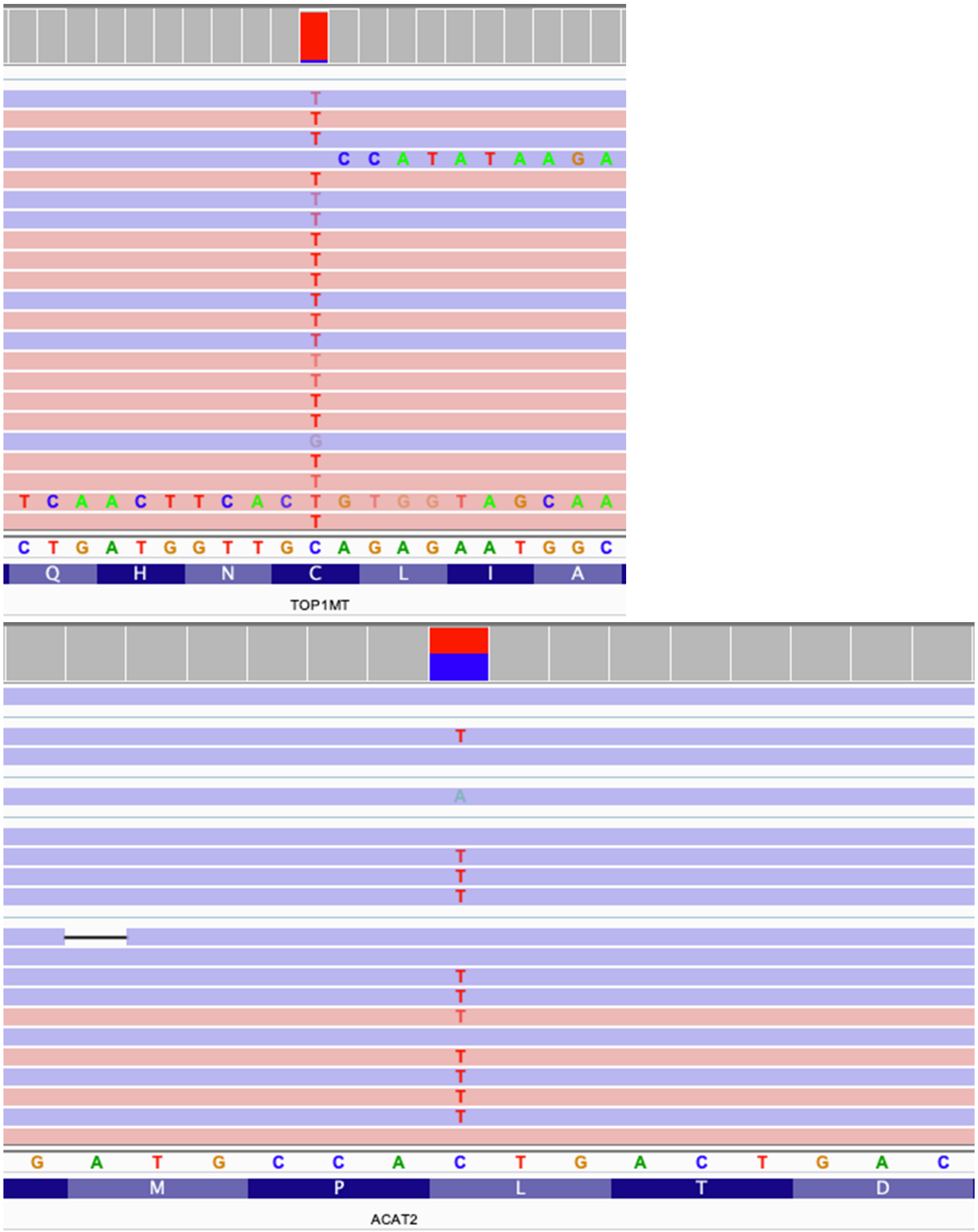

**Supplementary Figure S3. Single-cell level quantification of CRISPR induced indel per each guide-RNA.** The percentage of reads with CRISPR induced indels are quantified for each cell. For each target, indel percentages were calculated separately between cells with or without the presence of the target gRNA.

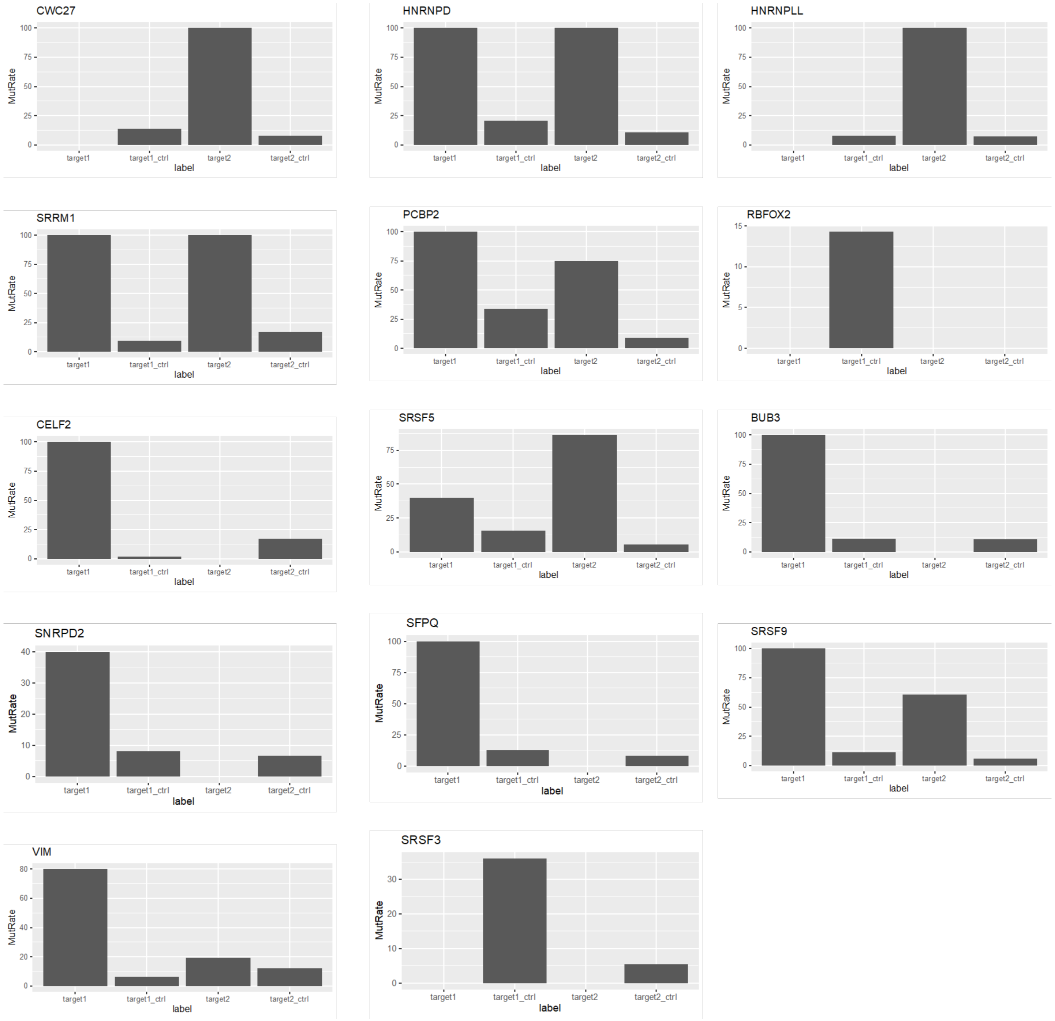

**Supplementary Figure S4. Single-cell level quantification of CRISPR induced exon skipping.** The upper panel shows the exon structure of *SRSF5* and how the gRNA affects exon4 and the resulting transcript structure. Red arrows indication PCR primers. CRISPR induced exon skipping events were quantified at single-cell resolution. Cells with gRNA targeting *SRSF5*'s exon4 had higher exon skipping frequency.

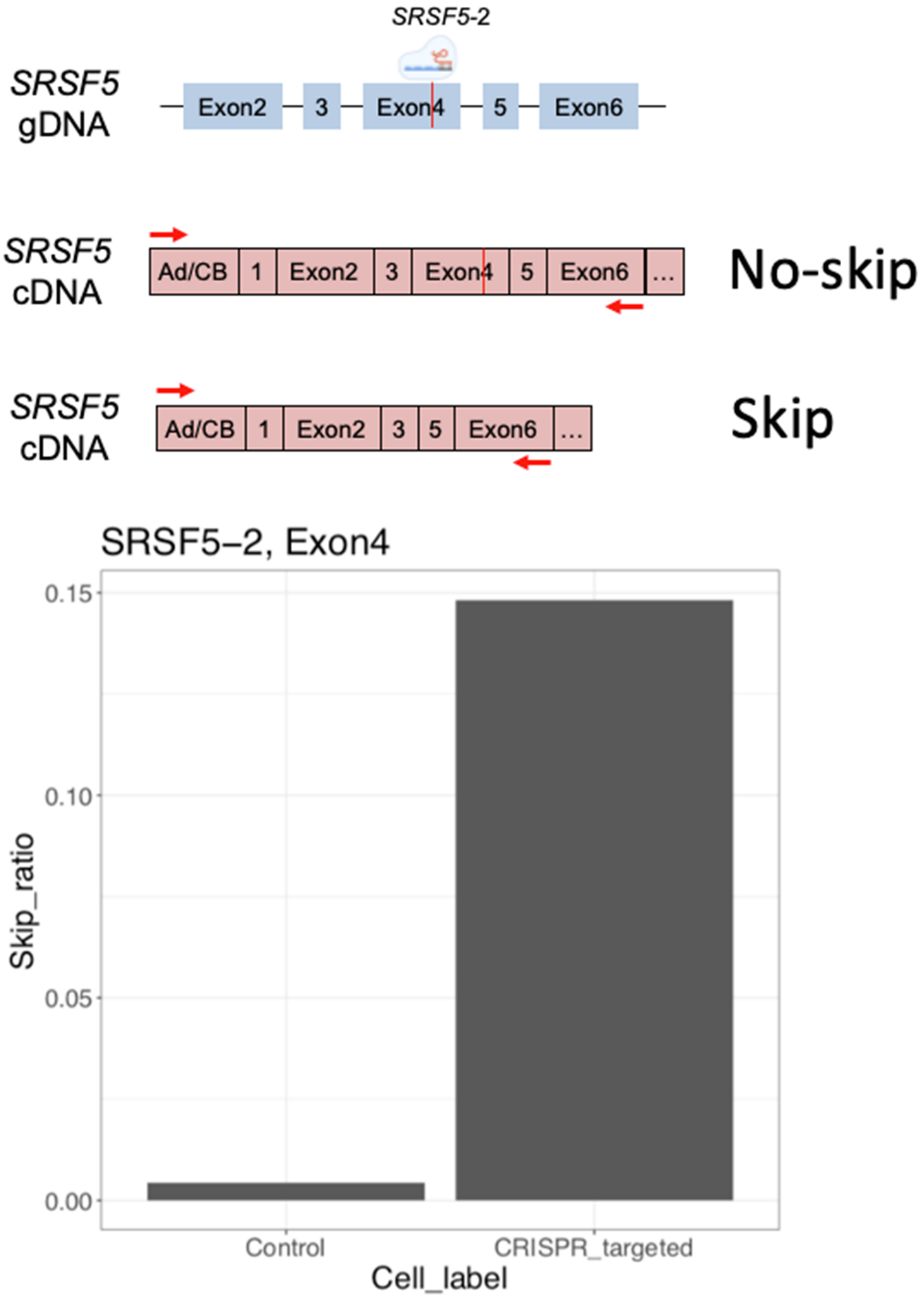

**Supplementary Figure S5. Amplicon versus adaptive sequencing omparison of single-cell level quantification of CRISPR induced indels.** CRISPR induced indel frequencies detected from amplicon sequencing in our previous study (x-axis) and adaptive sequencing in this study (y-axis) are compared.

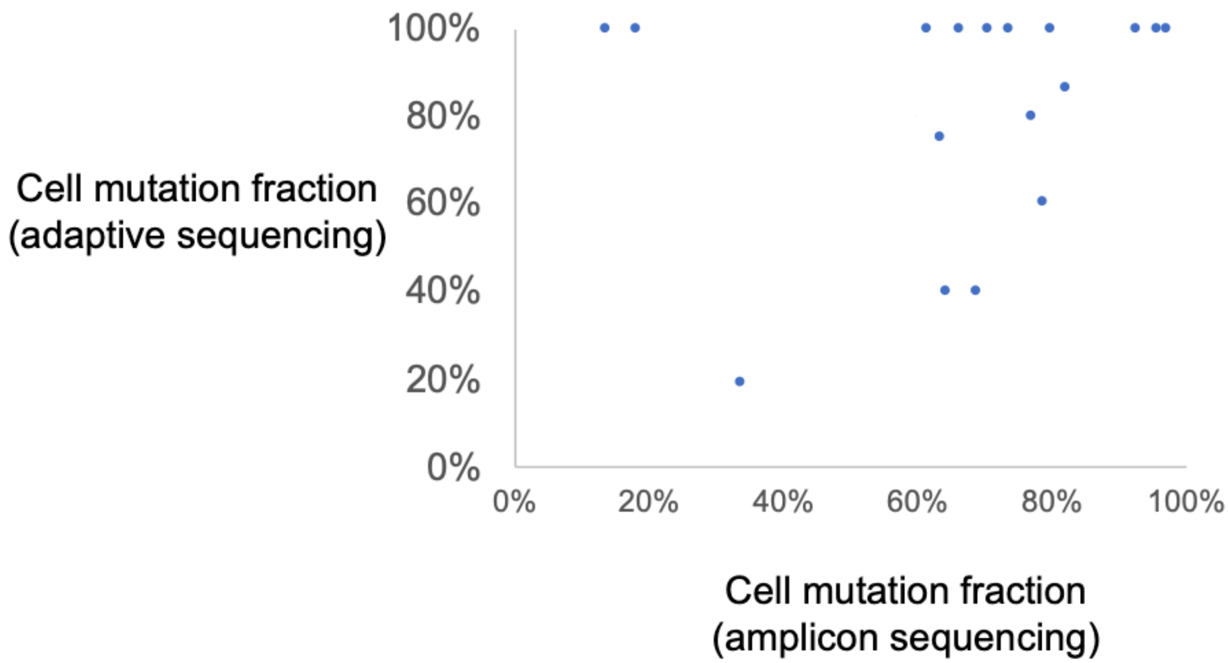

**Supplementary Figure S6. Comparison of long-read and short-read gene expression for patient tumor samples.** The heatmap shows the average UMI per cell/gene from short-reads for genes having substitution mutations. The scatter plots show the average UMI per cell from short-reads (x-axis) and total transcripts from long-reads (y-axis) for clinically mutated gene subset, and then all genes in gene panel. Scatter plots are very similar for both samples per patient so only one sample per patient is shown. **A.** T1/T2 average UMI; T1 scatter plots. **B.** T3/T4 average UMI; T3 scatter plots. **C.** T5/T6 average UMI; T5 scatter plots

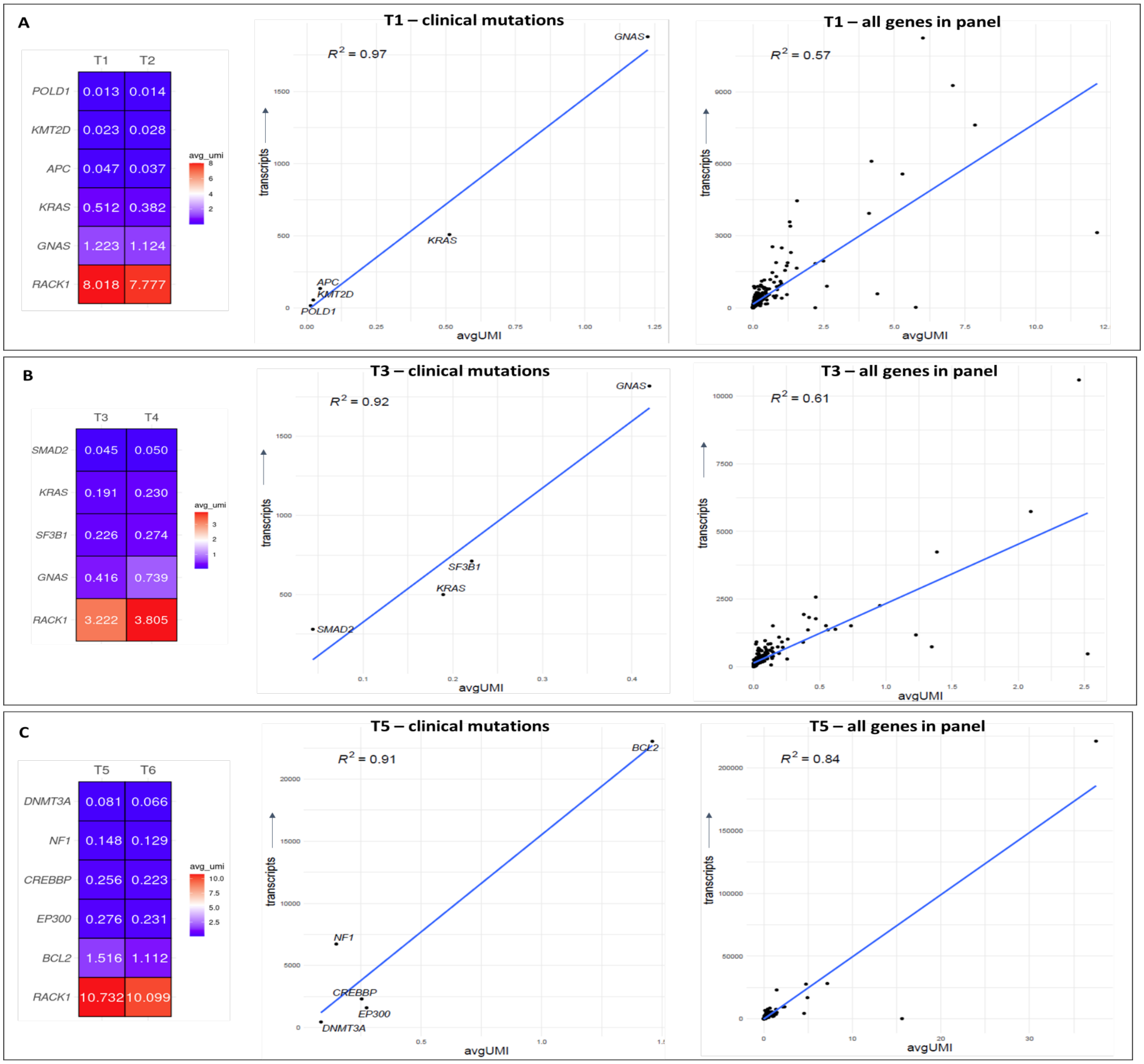
